# Regulation of Chromatin Accessibility by hypoxia and HIF

**DOI:** 10.1101/2022.01.07.475388

**Authors:** Michael Batie, Julianty Frost, Dilem Shakir, Sonia Rocha

## Abstract

Reduced oxygen availability (hypoxia) can act as a signalling cue in physiological processes such as development, but also in pathological conditions such as cancer or ischaemic disease. As such, understanding how cells and organisms respond to hypoxia is of great importance. The family of transcription factors called Hypoxia Inducible Factors (HIFs) coordinate a transcriptional programme required for survival and adaptation to hypoxia. The effects of hypoxia and HIF on the chromatin accessibility landscape are still unclear. Here, using genome wide mapping of chromatin accessibility via ATAC-seq, we find hypoxia induces loci specific changes in chromatin accessibility enriched at hypoxia transcriptionally responsive genes. These changes are predominantly HIF dependent, reversible upon reoxygenation and partially mimicked by chemical HIF stabilisation independent of molecular dioxygenase inhibition. This work demonstrates that indeed, HIF stabilisation is necessary and sufficient to alter chromatin accessibility in hypoxia, with implications for our understanding of gene expression regulation by hypoxia and HIF.

## Introduction

Molecular oxygen utilisation and sensing is an essential feature of metazoan life (1). Decreased oxygen availability (hypoxia) triggers a cellular response, central to which is the activation of transcriptional changes mediated by Hypoxia Inducible Family (HIF) transcription factors (2–5). HIF heterodimers, typically consisting of an oxygen labile α subunit (HIF-1α and HIF-2α), and a constitutively expressed β subunit (HIF-1β), bind DNA at hypoxia response elements (HREs), and typically function as gene transactivators (6, 7). Canonical regulation of HIF occurs via the Prolyl Hydroxylases (PHD)/ von Hippel–Lindau (VHL)/HIF axis. Under normal oxygen tensions, PHDs, a group of 2-OG dependent dioxygenases (2-OGDDs), proline hydroxylate HIF-1α and HIF-2α, targeting them for polyubiquitination by VHL E3 ubiquitin ligase complex and subsequent proteosomal degradation (1, 8). Impairment of PHDs activity in hypoxia, due to their oxygen dependence, stabilises HIF-α subunits and activates the HIF pathway.

At the chromatin level, HIF has been shown to predominantly bind RNA polII loaded, accessible chromatin regions with pre-established and primed, promoter enhancer loops (3, 5, 9, 10). HIF function is mediated by coactivators, including CREB-binding protein (CBP)/p300, SET Domain Containing 1B Histone Lysine Methyltransferase (SET1B), CDK8 and KAT5 (11). Chromatin also directly senses oxygen through 2-OGDDs (12–14). Inhibition of certain Ten-eleven Translocation (TET) methylcytosine dioxygenases and Jumonji C (JmjC)-domain containing histone demethylases in hypoxia alters DNA and histone methylation landscape respectively and coordinates transcriptional responses (12, 13, 15). Recently, several studies have used Assay for Transposase-Accessible Chromatin using sequencing (ATAC)-seq to explore the chromatin accessibility landscape in response to oxygen fluctuation (16–19). These studies reveal that hypoxia induces dynamic changes in chromatin accessibility in cell culture models. However, the roles of HIF and 2-OGDD oxygen sensing in hypoxia induced chromatin accessibility have not been studied, and remains an important question, as more inhibitors of these pathways are developed to enter the clinical setting.

Here, using ATAC-seq, we have investigated effects of oxygen deprivation and reoxygenation on chromatin accessibility in cells in culture. We also measured transcript changes in hypoxia with RNA-seq and analysed the role of HIF in this process using a specific stabiliser of HIF-α as well as siRNA-mediated depletion of HIF-1β. Integrative analysis of ATAC-seq with RNA-seq reveals that hypoxia induces coordinated and specific changes to chromatin accessible regions, which correlate with gene expression changes. Furthermore, most hypoxia inducible changes to chromatin accessibility are HIF dependent and rapidly reversible upon reoxygenation. Additionally, HIF stabilisation, independent of 2-OGDD inhibition, is sufficient to partially mimic hypoxia-induced changes in chromatin accessibility. HIF binding sites are also enriched at genomic loci with hypoxia inducible increases in chromatin accessibility. This demonstrates a central role for HIF in controlling chromatin accessibility dynamics in response to hypoxia. Lastly, we find that H3K4me3 levels correlate with accessibility changes in hypoxia and provide evidence for a role of KDM5A in regulation of chromatin accessibility in hypoxia.

## Results

### Genome wide mapping of the chromatin accessibility landscape in normoxia and hypoxia

The hypoxia response in cells involves a coordinated transcriptional programme (6). However, chromatin accessibility dynamics in response to hypoxia are not well defined. To investigate the effect of acute hypoxia on chromatin accessibility, we performed ATAC-seq in HeLa cells cultured at 21% oxygen (control) or exposed to 1 and 24h of hypoxia (1% oxygen) (Figure 1A, Supplementary Dataset 1). 71,651 high confidence (identified in all biological replicates within a condition, each with an FDR < 1×10^-15^) open chromatin regions (ORs) are identified across all time points, with 80% present in all time points (Figure 1A). 23,901 OR genes (ORs at genic regions) are identified across all time points, with 91% present in all time points (Supplementary Figure S1A). Data is in concordance with current ENCODE standards for ATAC-seq (Supplementary Dataset 1) and similar regions of open chromatin are identified comparing to other published HeLa ATAC-seq (Supplementary Figure S1B), demonstrating high data quality.

**Figure 1.**
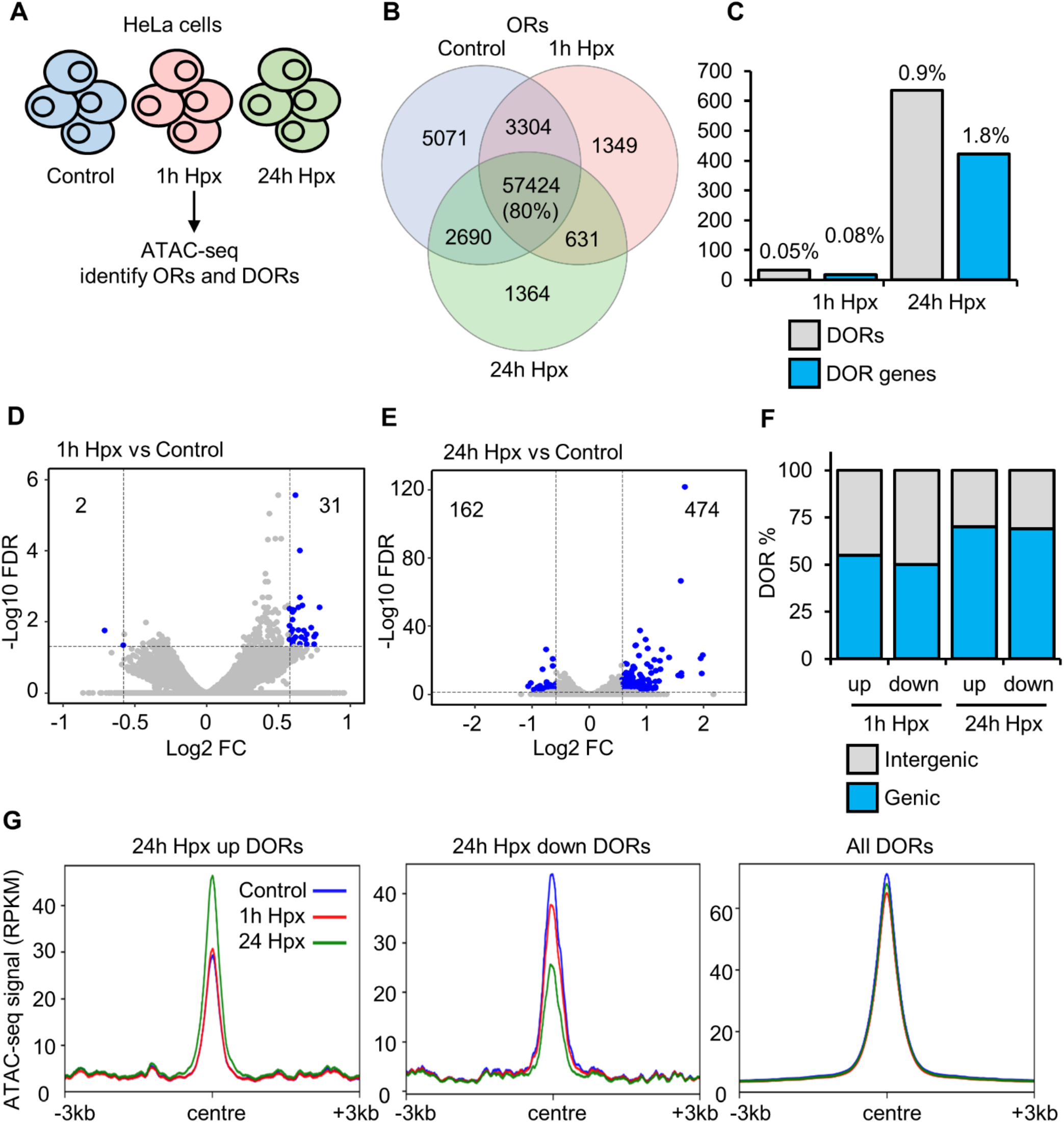
Chromatin accessibility changes in response to hypoxia. **A)** ATAC-seq in HeLa cells cultured at 21% oxygen, transfected with control siRNA and exposed to 0h (control), 1h and 24h 1% oxygen (hypoxia (Hpx)). **B)** Overlap of open chromatin regions (ORs). **C)** Number of high stringency (log2 fold change -/+0.58 and FDR <0.05) differentially open chromatin regions (DORs) and genes with DORs (DOR genes), and percentage relative to total ORs and OR genes. **D-E)** Volcano plots for 1h Hypoxia vs control DOR analysis and 24h hypoxia vs control DOR analysis (s=significant, ns= non-significant). **F)** Genomic location of DORs. **G)** Metagene plots of ATAC-seq signal (RPKM) at the indicated regions.

Differential open region (DOR) analysis (Supplementary Dataset 2) identified site specific changes at ORs in response to hypoxia. 33 high stringency (log2 fold change -/+0.58 and FDR <0.05) DORs are present in response to 1h hypoxia (Figure 1C, D) and 636 high stringency DORs are present in response to 24h hypoxia (Figure 1C, E). 18 DOR genes and 422 DOR genes are identified in response to 1h and 24h hypoxia respectively (Figure 1C). Of the 1h hypoxia DORs, 31/33 are upregulated and 2/33 are downregulated (Figure 1D). When analysing the 24h hypoxia DORs, 474/636 are upregulated and 162/636 are downregulated (Figure 1E). DORs are spread across genic and intergenic regions (Figure 1F). Mapping ATAC-seq signal across hypoxia DORs shows changes in chromatin accessibility induced by hypoxia are loci specific (Figure 1G, Supplementary Figure S1C-D). When using lower stringency DOR analysis (FDR<0.1) we find 445 DORs and 336 DOR genes in response to 1h hypoxia, and 4877 DORs and 2955 DOR genes in response to 24h hypoxia (Supplementary Figure S1E-F). Interestingly, this lower stringency analysis produced similar number of changes to those identified in HUVEC cells exposed to hypoxia (16).

These results show that hypoxia induces changes in chromatin accessibility at a specific set of loci in HeLa cells, with most changes occurring at later than 1h of hypoxia.

**Supplementary Figure S1.**
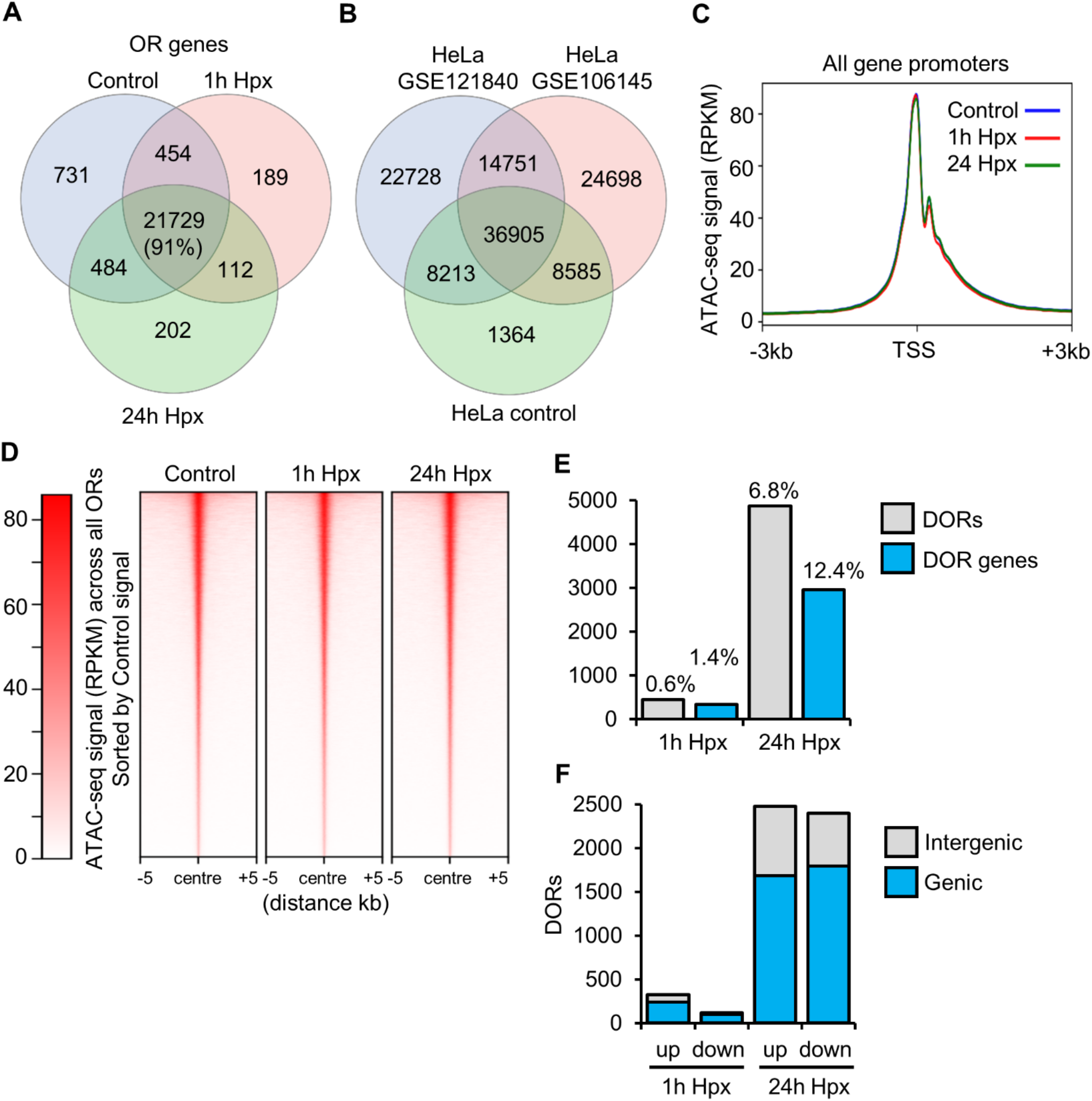
Chromatin accessibility changes in response to hypoxia additional data. ATAC-seq in HeLa cells cultured at 21% oxygen, transfected with control siRNA and exposed to 0h (control), 1h and 24h 1% oxygen (hypoxia (Hpx)). **A)** Overlap of open chromatin region (OR) genes. **B)** Overlap of ORs between different HeLa ATAC-seq studies. **C)** Metagene plots of ATAC-seq signal (RPKM) at all gene promoters. **D)** Heatmap of ATAC-seq signal across all ORs and ranked by control high to low signal ORs. **E)** Number of low stringency (FDR <0.1) differentially open chromatin regions (DORs) and genes with DORs (DOR genes), and percentage relative to total ORs and OR genes. **F)** Number of upregulated and downregulated ORs and their genomic location.

### Hypoxia induced changes in chromatin accessibility correlate with changes in gene expression

Hypoxia responsive chromatin accessible regions were investigated for the associated gene signatures (Figure 2A). Glycolysis, hypoxia and EMT gene signatures are enriched at 24h hypoxia upregulated DOR genes (Figure 2B). No statistically significant pathway enrichment was found for 24h hypoxia downregulated DOR genes or 1h hypoxia DOR genes. To determine how changes in gene expression correlate with changes in chromatin accessibility, we performed RNA-seq in HeLa cells exposed to 0, 1 and 24h of hypoxia (Supplementary Dataset 3). 25 (23 upregulated, 2 downregulated) differentially expressed genes (DEGs) in response to 1h hypoxia (Supplementary Dataset 3), and 1330 DEGs (1088 upregulated, 242 downregulated) in response to 24h hypoxia are identified from this analysis (Supplementary dataset 3). From integrative analysis of ATAC-seq with RNA-seq data, we found a subset of genes with hypoxia induced differential expression, which also possess changes in chromatin accessibility (Figure 2C-D; Supplementary Dataset 4). 24h hypoxia upregulated DOR genes show significant correlation with 24h hypoxia upregulated expression genes, 37 of the 24h hypoxia upregulated DOR genes have increased gene expression (Figure 2C), among these are the well characterised, core hypoxia responsive genes, *CA9, NDRG1* and *EGNL3* (protein name PHD3) (Figure 2D). GeneSet Enrichment Analysis also confirmed these results (Figure 2E). Interestingly, 24h hypoxia downregulated OR genes also show significant correlation with 24h hypoxia downregulated expression genes (Figure 2C-D).

**Figure 2.**
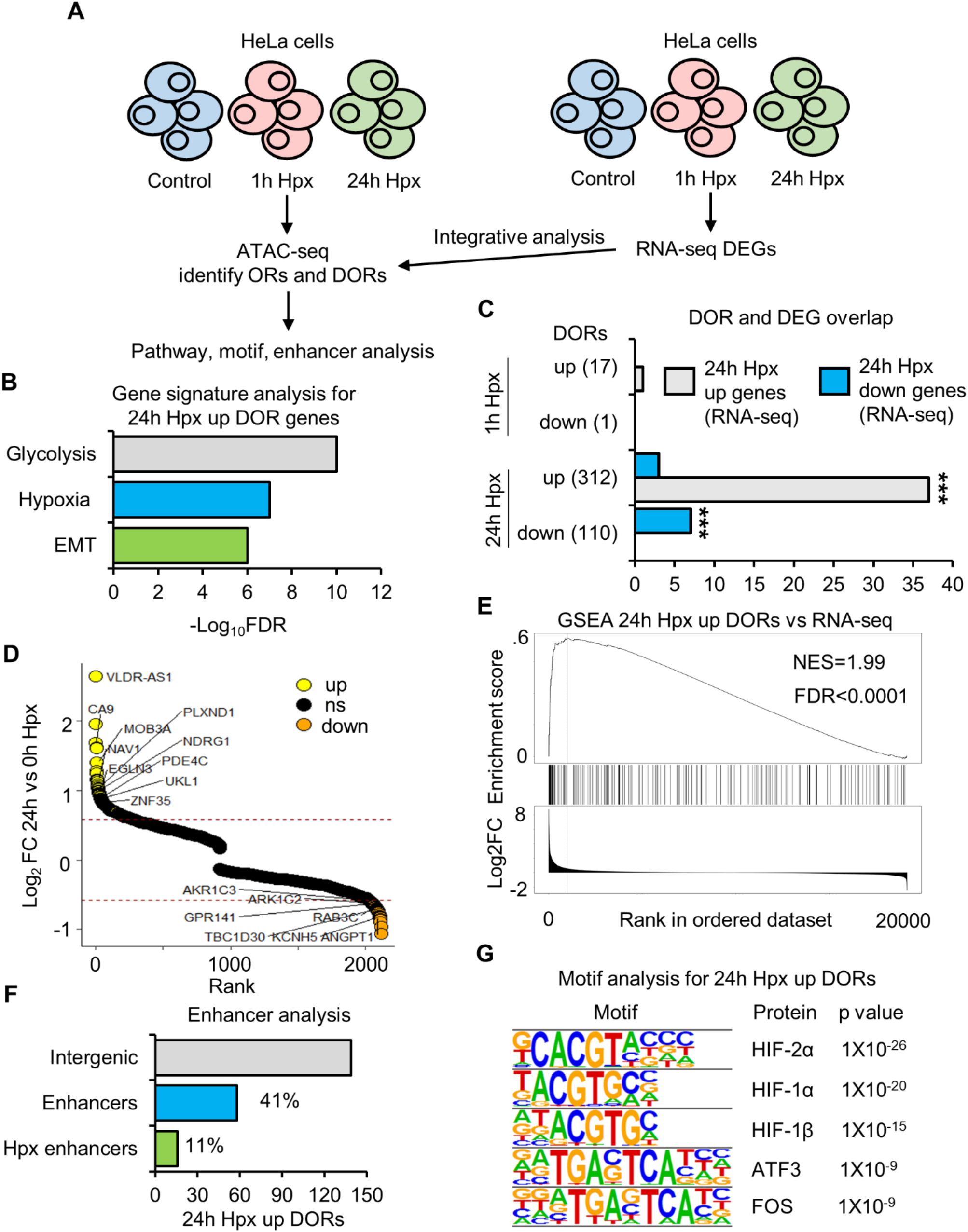
Hypoxia inducible changes in open chromatin are enriched at hypoxia transcriptionally regulated genes. **A)** ATAC-seq and RNA-seq in HeLa cells cultured at 21% oxygen, transfected with control siRNA and exposed to 0h (control), 1h and 24h 1% oxygen (hypoxia (Hpx)). **B)** Gene signature analysis. **C)** Overlap between genes with differentially open chromatin regions (DOR genes) and genes with differential RNA expression (DEG) in response to hypoxia (\*\*\**P*< 0.001). **D)** Gene list ranked from high to low fold change in chromatin accessibility in response to 24h hypoxia. Some hypoxia upregulated DEG and DOR genes and downregulated DEG and DOR genes are labelled. **E)** GeneSet Enrichment Analysis between 24h hypoxia upregulated DOR genes and a list of genes ranked from high to low 24h hypoxia RNA expression fold change. **F)** Percentage of 24h hypoxia upregulated DORs at intergenic regions that are active enhancers and active enhancers linked to the promoters of genes with 24h hypoxia upregulated RNA expression. **G)** Motif enrichment analysis.

The aforementioned analysis is specific to genic (promoter and gene body) DORs. To functionally annotate changes in chromatin accessibility at intergenic regions, we performed enhancer analysis and found that 42% of intergenic 24h hypoxia upregulated DORs are at active enhancers (Figure 2E). 11% (16/142) are enhancer partners for the promoters of genes whose expression is upregulated at 24h hypoxia (Figure 2F). These include the promoters of the well characterised, core hypoxia responsive genes, *SCL2A3* (protein name GLUT3) and *NDRG1*. Thus, changes in accessibility at hypoxia responsive genes occur at both gene proximal and distal regulatory elements. Lastly, HIF subunits motifs are enriched at 24h hypoxia upregulated DORs (Figure 2G), suggesting a role of HIF in coordination of changes in chromatin accessibility in hypoxia.

### Hypoxia induced changes in chromatin accessibility are mostly sensitive to reoxygenation and are HIF dependent

HIF is the master regulator of transcriptional changes in response to hypoxia; however, its role in chromatin accessibility regulation has been elusive. To ascertain the dependence of HIF on hypoxia inducible changes to chromatin accessibility, we performed ATAC-seq in cells where HIF-1β (the obligate partner for HIF heterodimer complexes) was depleted by siRNA prior to hypoxia exposure (Figure 3A, Supplementary Dataset 1, 2, 5). 1h hypoxia DORs were almost exclusively dependent on HIF-1β (Figure 3B, Supplementary Dataset 5). At 24h hypoxia, requirement for HIF is favoured at sites with increased accessibility over reduced accessibility, with 92% of 24h hypoxia upregulated ORs and 57% of 24h hypoxia downregulated DORs requiring HIF-1β (Figure 3B, Supplementary Dataset 5). Hypoxia DORs are classed as dependent on HIF-1β if they are they not identified as DORs with HIF-1β siRNA treatment when comparing to control (0h hypoxia, control siRNA). Validation of siRNA depletion of HIF-1β is confirmed by immunoblotting (Supplementary Figure S2A).

**Figure 3.**
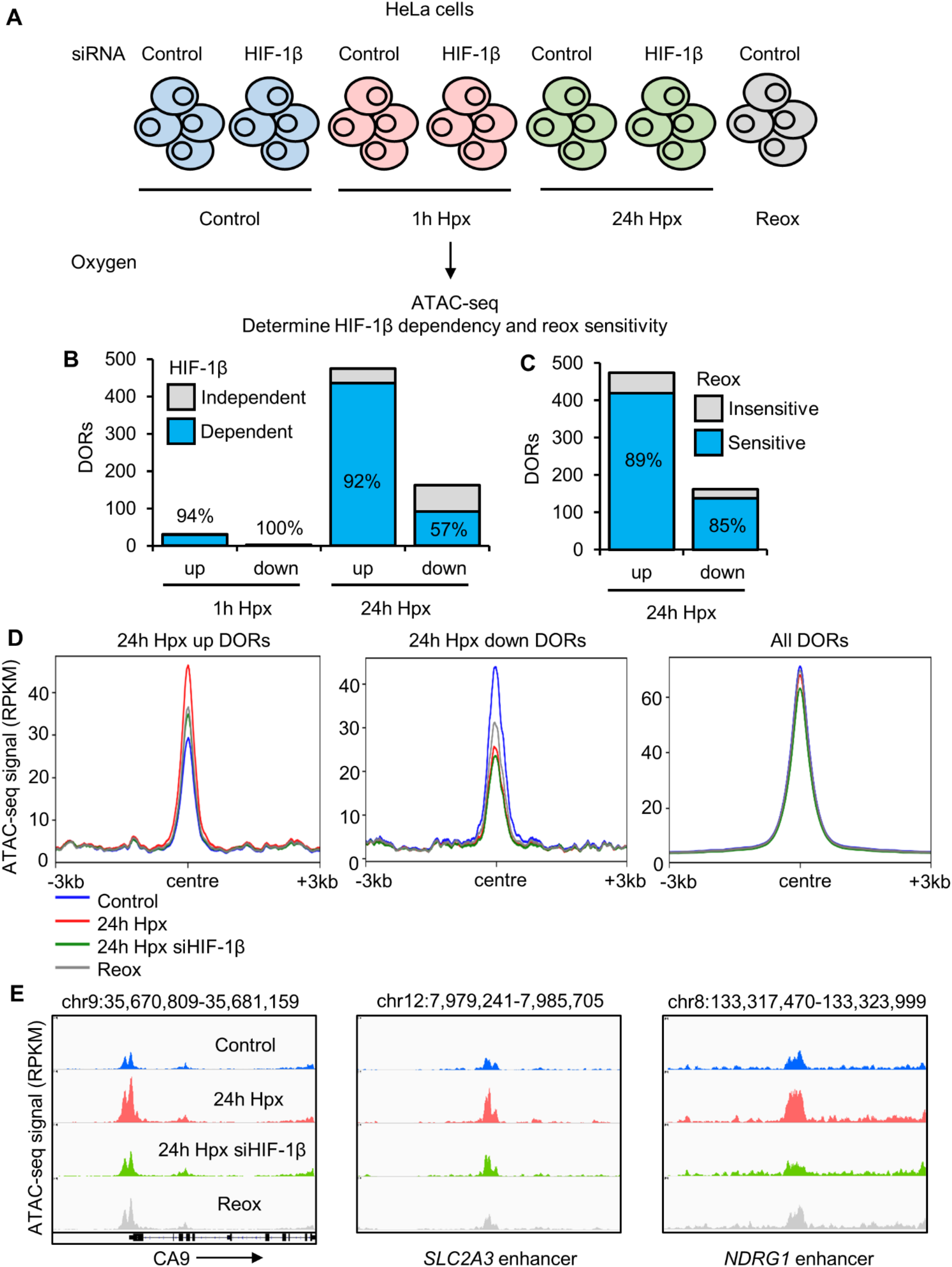
Hypoxia inducible changes in open chromatin are mainly sensitive to reoxygenation HIF dependent. **A)** ATAC-seq in HeLa cells cultured in 21% oxygen; transfected with control siRNA or HIF-1β siRNA, and exposed to 0h (control), 1h, 24h 1% oxygen (hypoxia (Hpx)) and 24h hypoxia followed by 1h at 21% oxygen (reoxygenation (Reox)). **B)** HIF-1β dependence of hypoxia differentially open chromatin regions (DORs), percentage of HIF-1β dependent DORs are labelled. **C)** Reoxygenation sensitivity of 24h hypoxia DORs, percentage of reoxygenation sensitive DORs are labelled. **D)** Metagene plots of ATAC-seq signal (RPKM) at the indicated regions. **E)** Coverage tracks of ATAC-seq signal at the *CA9* promoter, *SLC2A3* (protein name GLUT3) enhancer and *NDRG1* enhancer.

To elucidate the sensitivity of hypoxia induced changes in chromatin accessibility to fluctuations in oxygen levels, we included a reoxygenation condition in our analysis (24h hypoxia (1% oxygen), followed by 1h at normoxia (21%> oxygen) (Figure 3A, Supplementary Dataset 1, 2, 6). The vast majority of 24h hypoxia DORs (89% of upregulated DORs and 85% of downregulated DORs) return to near normoxic levels upon reoxygenation (Figure 3B, Supplementary Dataset 6). DORs are classed as reoxygenation sensitive if they are not identified as DORs in reoxygenation condition compared to control (0h hypoxia, control siRNA). As a control for hypoxia and reoxygenation, immublotting of HIF-1α was performed (Supplementary Figure S2B). As expected, HIF-1α increases at 1h and 24h hypoxia and this increase is lost with reoxygenation.

PCA analysis shows ATAC-seq sample clustering by treatment (Supplementary Figure S2C). As found with the analysis of hypoxia treatment, reoxygenation and HIF-1β depletion cause loci specific changes as opposed to genome wide changes in chromatin accessibility (Figure 3D, Supplementary Figure S2D-E). Coverage tracks of a subset of hypoxia upregulated DORs at hypoxia transcriptionally upregulated gene promoters/enhancers are displayed, demonstrating HIF-1β dependence and reoxygenation sensitivity of hypoxia inducible chromatin accessibility changes.

These data indicate that changes in chromatin accessibility in hypoxia are highly dependent on HIF, particularly regards to loci with increased accessibility in hypoxia. In addition, hypoxia-induced chromatin changes are dependent on oxygen availability and rapidly reversed upon re-oxygenation.

**Supplementary Figure S2.**
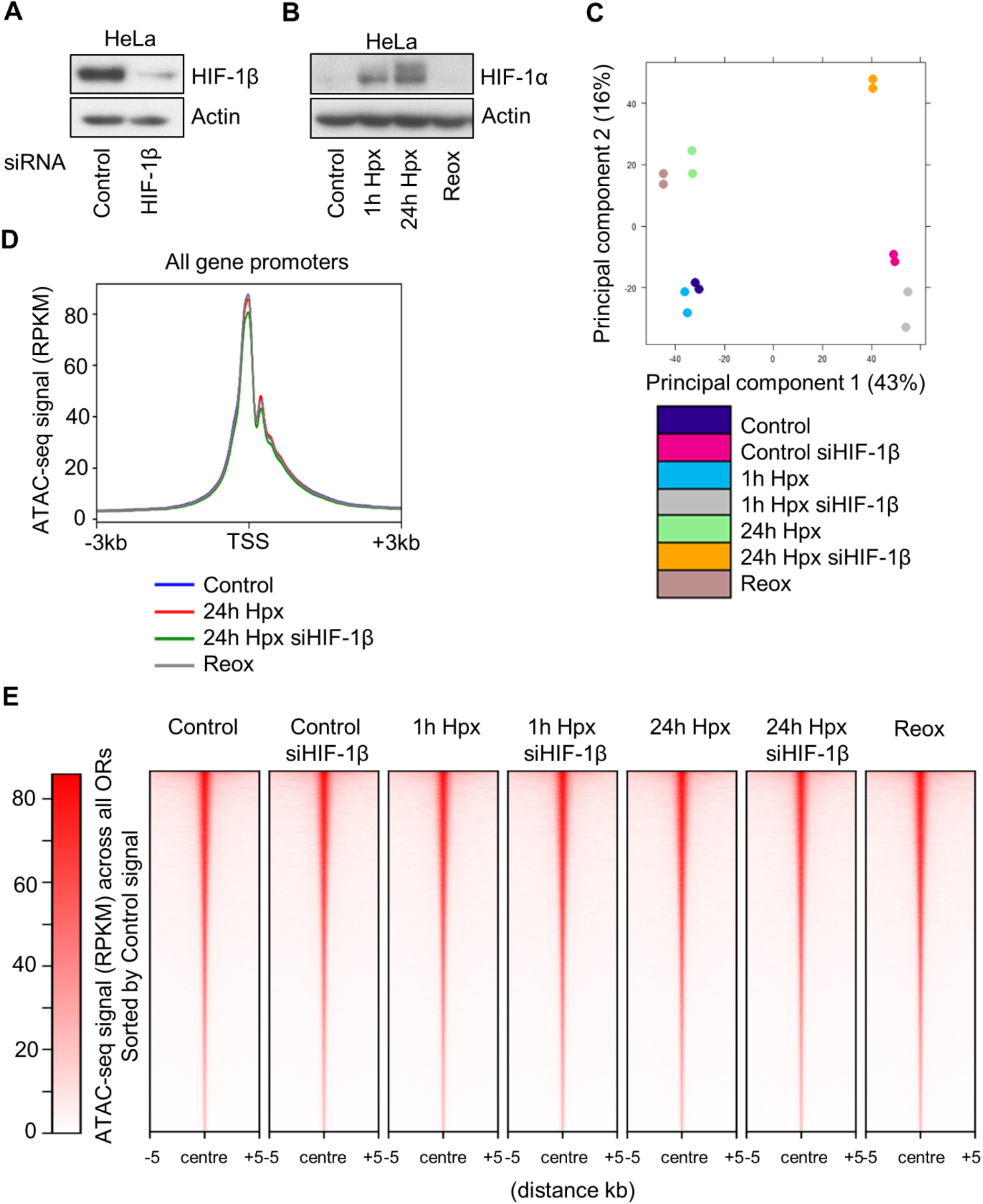
Hypoxia inducible changes in open chromatin are mainly sensitive to reoxygenation HIF dependent additional data. HeLa cells were cultured in 21% oxygen; transfected with control siRNA or HIF-1β siRNA, and exposed to 0h (control), 1h, 24h 1% oxygen (hypoxia (Hpx)) and 24h hypoxia followed by 1h at 21% oxygen (reoxygenation (Reox)). **A, B)** Immunoblot of the indicated proteins. **C)** ATAC-seq principal component analysis (PCA). **D)** Metagene plots of ATAC-seq signal (RPKM) at all gene promoters. **E)** Heatmap of ATAC-seq signal across all ORs and ranked by control high to low signal ORs.

### VH298 mediated HIF stabilisation is sufficient to induce changes to chromatin accessibility

In addition to HIF stabilisation, 2-OGDD inhibition in hypoxia can alter modifications of other targets, including histones (12, 13). To uncouple the effects of HIF stabilisation and 2-OGDD inhibition on chromatin accessibility, we performed ATAC-seq in HeLa cells treated for 24h with DMSO (control) and 24h, 100 μM VH298, a specific chemical inhibitor of the hydroxylated HIF-α binding pocket of VHL (20–22) (Figure 4A, Supplementary Dataset 1). Immunoblot analysis confirmed HIF-1α stabilisation in response to VH298 treatment (Supplementary Figure S3A). 72,137 high confidence (identified in all biological replicates within a condition, each with an FDR < 1×10^-15^) ORs are found across control and VH298 treated samples, with 85% found in both (Supplementary Figure S3B). 23,993 OR genes are present across control and VH298 treated samples, with 93% present in both (Supplementary Figure S3B). VH298 DOR analysis reveals 447 high stringency (log2 fold change -/+0.58 and FDR <0.05) DORs and 292 DOR genes in response to 24h VH298 treatment (Supplementary Figure S3C, Supplementary Dataset 2). Of the VH298 DORs, 318/447 are upregulated and 129/447 are downregulated. (Figure 4B). 67% of upregulated VH298 DORs and 72% of VH298 downregulated DORs are at genic regions (Supplementary Figure S3D). As with hypoxia, VH298 induces loci specific changes in chromatin accessibility (Supplementary Figure S3E-F). Lower stringency DOR analysis (FDR <0.1) finds 1555 DORs and 1031 DOR genes in response to VH298 treatment (Supplementary Figure S3G-H).

**Figure 4.**
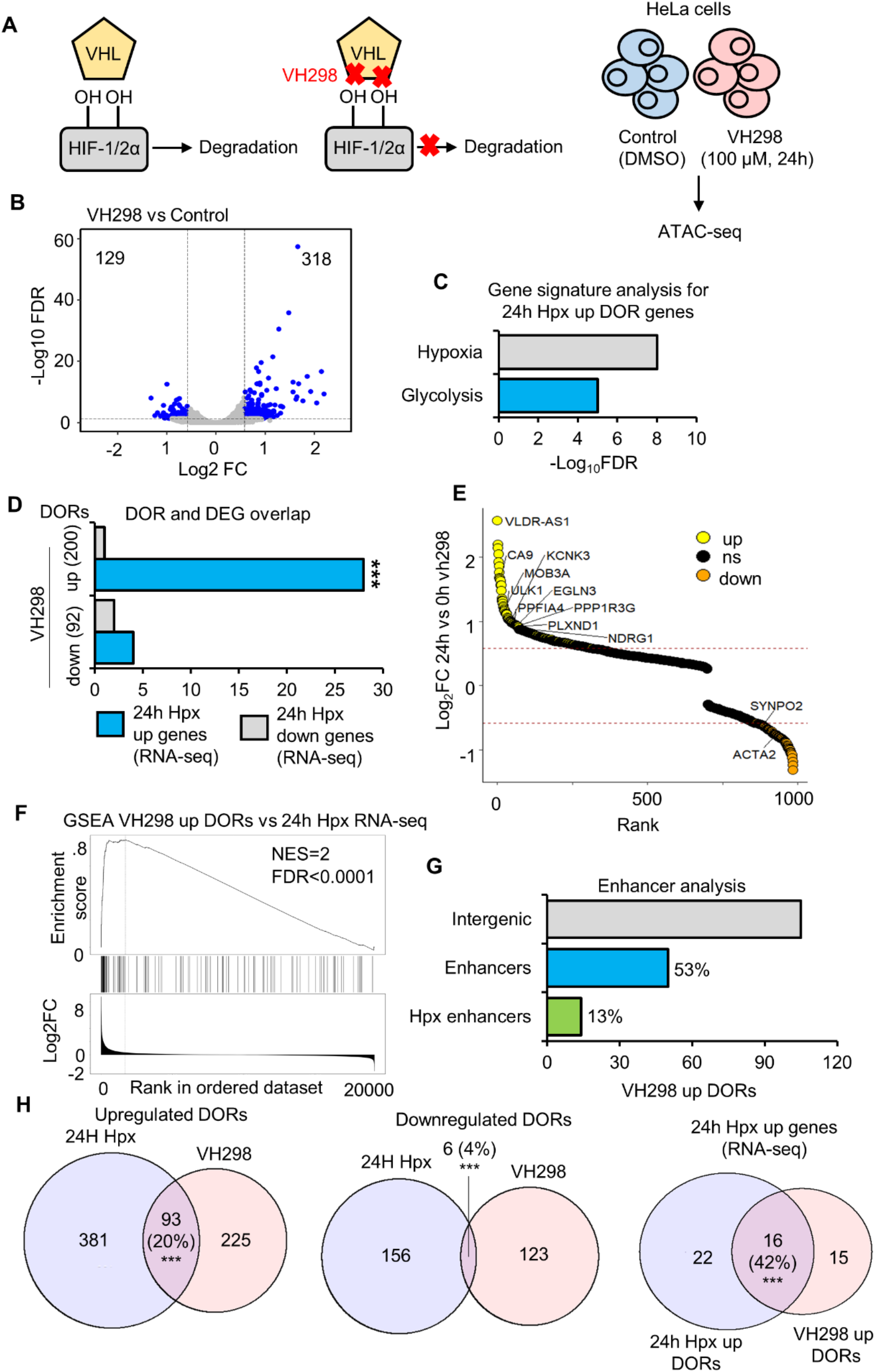
Chromatin accessibility changes in response to HIF stabilisation via VH298. **A)** ATAC-seq in HeLa cells cultured at 21% oxygen and treated with DMSO (control) and 100 μM VH298 for 24h. **B)** Volcano plot for differentially open chromatin region (DOR) analysis (s=significant, ns= non-significant). **C)** Gene signature analysis. **D)** Overlap between genes with differentially open chromatin regions (DOR genes) in response to VH298 and genes with differential RNA expression (DEGs) in response to hypoxia (\*\*\**P*< 0.001). **E)** Gene list ranked from high to low fold change in chromatin accessibility in response to VH298. Some upregulated hypoxia DEG and VH298 DOR genes and downregulated hypoxia DEG and VH298 DOR genes are labelled. **F)** GeneSet Enrichment Analysis between VH298 upregulated OR genes and a list of genes ranked from high to low 24h hypoxia RNA expression fold change. **G)** Percentage of VH298 upregulated DORs at intergenic regions that are active enhancers and active enhancers linked to the promoters of genes with 24h hypoxia upregulated RNA expression. **H)** Overlap of VH298 and 24h hypoxia DORs, \*\*\**P*< 0.001.

Gene signature analysis identified hypoxia and glycolysis pathways as enriched at VH298 upregulated DOR genes (Figure 4C). These signatures were also enriched in the data related to 24h hypoxia. Similarly, as with 24h hypoxia exposure, VH298 upregulated DOR genes have significant overlap with 24h hypoxia upregulated expression genes (Figure 4D-E), and are enriched at 24h hypoxia upregulated expression genes as determined by GeneSet Enrichment Analysis (Figure 4F). Integrative analysis with the HACER enhancer database shows that 53% of intergenic VH298 upregulated DORs are at enhancer regions and 13% (14/105) are enhancer partners linked to promoters of 24h hypoxia upregulated expression genes (Figure 4G). Motif enrichment analysis reveals HIF subunit-binding motifs are overrepresented in VH298 upregulated DOR genes (Supplementary Figure S3I). These data show that, HIF stabilisation, independent of dioxygenase inhibition, is sufficient to trigger loci specific changes in chromatin accessibility linked to hypoxia regulated genes.

We next directly compared chromatin accessibility responses between hypoxia and VH298 (Supplementary Dataset 7). Exposure to 24h hypoxia induced changes in chromatin accessibility at 42% more genomic loci than 24h VH298 treatment (636 DORs compared to 447 DORs). There is higher similarity of upregulated responses, with 20% of hypoxia upregulated sites also upregulated by VH298 whereas only 4% of hypoxia downregulated sites are also downregulated by VH298 (Figure 4H). A greater correlation between hypoxia and VH298 accessibility changes is observed when comparing changes located at hypoxia upregulated expression genes (Figure 4H). 53 hypoxia upregulated expression genes (RNA-seq) display increased accessibility in response to hypoxia or VH298, sharing 16 gene regions, 22 unique to hypoxia treatment and 15 unique to VH298 treatment (Figure 4H).

This analysis establishes that VH298 partially mimics the hypoxia response, concerning loci specific increases in accessibility. Thus, HIF stabilisation, independent of oxygen sensing enzyme inhibition, is sufficient to drive a subset of hypoxia inducible changes in chromatin accessibility.

**Supplementary Figure S3.**
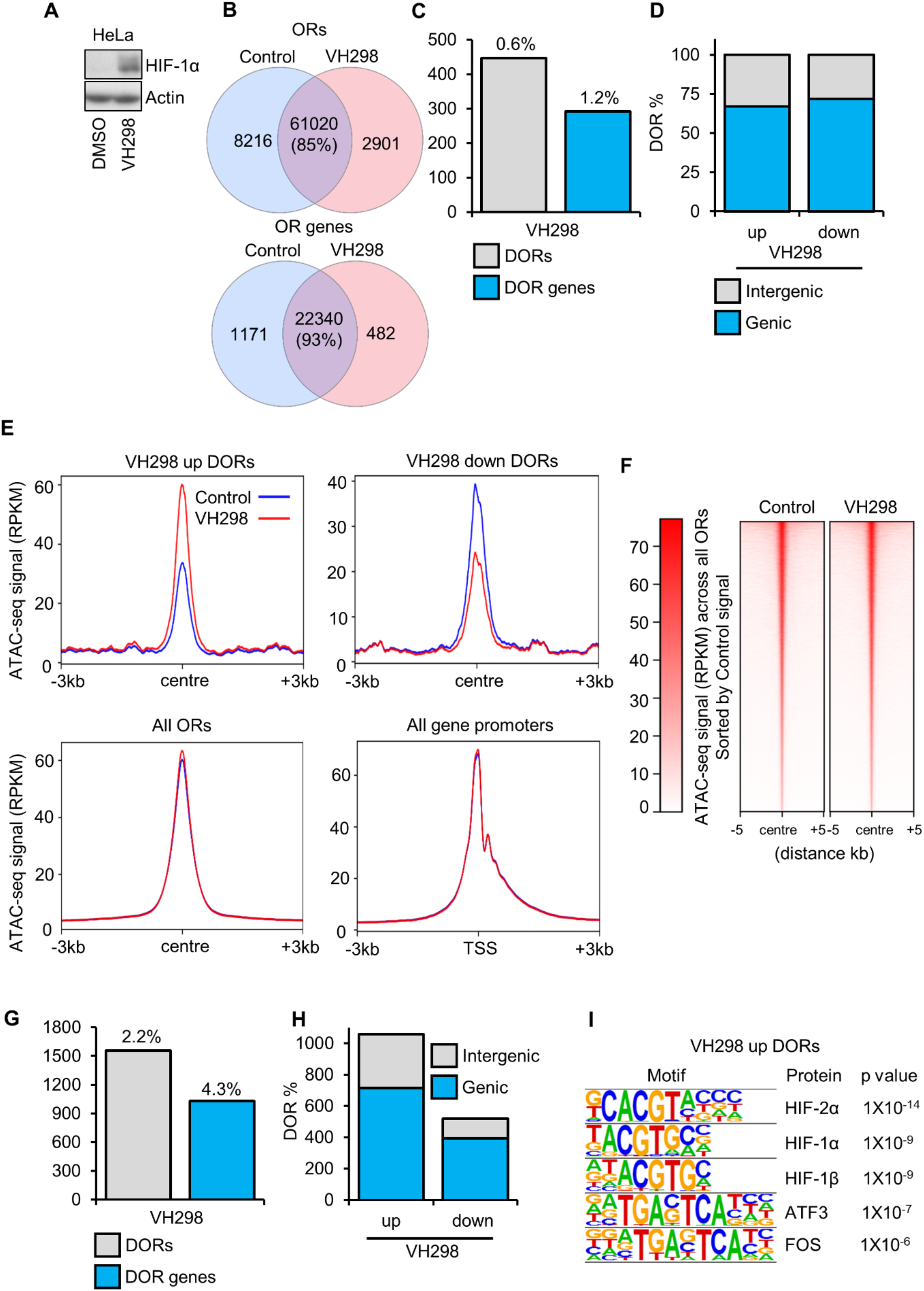
VH298 ATAC-seq additional data. HeLa cells were cultured at 21% oxygen and treated with DMSO (control) and 100 μM VH298 for 24h. **A)** Immunoblot of the indicated proteins. **B)** ATAC-seq overlap analysis of open chromatin regions (ORs) and OR genes. **C)** ATAC-seq analysis, number of high stringency (log2 fold change -/+0.58 and FDR <0.05) differentially open chromatin regions (DORs) and genes with DORs (DOR genes), and percentage relative to total ORs and OR genes. **D)** ATAC-seq analysis, number of upregulated and downregulated DORs and their genomic location. **E)** Metagene plots of ATAC-seq signal (RPKM) plotted indicated DORs and protein coding gene promoters. **F)** Heatmap of ATAC-seq signal, ranked by control high to low signal ORs. **G)** ATAC-seq analysis, number of low stringency (FDR <0.1) differentially open chromatin regions (DORs) and genes with DORs (DOR genes), and percentage relative to total ORs and OR genes. **H)** ATAC-seq analysis, number of upregulated and downregulated DORs (low stringency DORs) and their genomic location. **I)** ATAC-seq motif enrichment analysis at VH298 upregulated DORs (high stringency).

### Hypoxia and VH298 induced accessibility changes are also observed by ATAC-qPCR

To confirm changes in chromatin accessibility in response to hypoxia and VH298 treatment, ATAC-qPCR analysis was performed on a set of loci identified by the ATAC-seq analysis (Figure 5). The *CA9* promoter displayed increased accessibility in response to 24h hypoxia exposure and 24h VH298 treatment, and this increase was reduced when HIF-1β is depleted in 24h hypoxia exposed cells and when cells are reoxygenated following 24h hypoxia exposure (Figure 5A-B). These results agree with the ATAC-seq analysis. Similar results were obtained for the *EGLN3* (protein name PHD3) gene body, *VLDLR AS-1* promoter, *NDRG1* enhancer and *SLC2A3* (protein name GLUT3) enhancer loci (Figure 5A-B). Also agreeing with the ATAC-seq analysis, promoter chromatin accessibility at the *FGF11* promoter was specifically increased in response to 24h hypoxia but not 24h VH298 treatment (Supplementary Figure S4A-B). To determine if these changes are also present in another human cancer cell line, we repeated 24h hypoxia and VH298 treatment ATAC-qPCR analysis in A549 cells (Figure 5C). Increased accessibility in response to 24h hypoxia and VH298 treatment at *EGLN3, VLDLR AS-1, NDRG1 and SLC2A3* was also found in A549 cells, although the increase in *SLC2A3* was not statistically significant (Figure 5C). No significant changes were present at the *CA9* promoter loci (Figure 5C). *FGF11* accessibility was also unaffected by VH298 treatment in both cell lines, and hypoxia upregulated accessibility was only observed in HeLa cells (Supplementary Figure S4C). Absence of increased accessibility in response to hypoxia at *CA9* and *FGF11* loci in A549 cells is not explained by lack of transcript upregulation, as both genes are upregulated in response to 24h hypoxia (determined by A549 RNA-seq (Supplementary Dataset 3), and could be due to a difference in timing of chromatin changes or represent cell type heterogeneity in hypoxia inducible chromatin accessibility changes.

**Figure 5.**
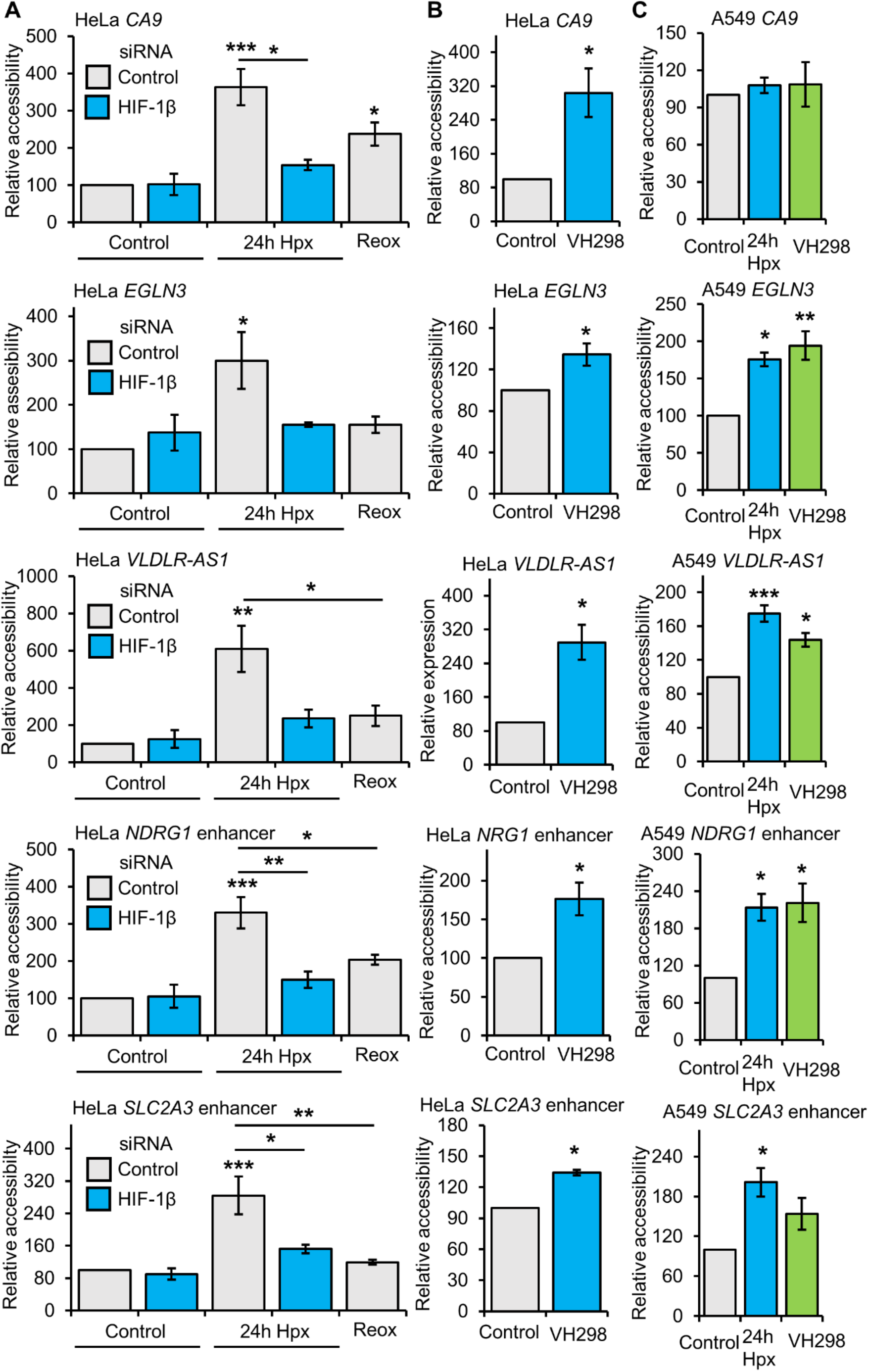
Validation of accessibility changes. **A)** ATAC-qPCR analysis in HeLa cells cultured at 21% oxygen, transfected with control siRNA or HIF-1β siRNA, and exposed to 0h (control), 24h 1% oxygen (hypoxia (Hpx)) and 24h hypoxia followed by 1h at 21% oxygen (reoxygenation). **B)** ATAC-qPCR analysis in HeLa cells cultured at 21% oxygen and treated with 24h DMSO (control) or 24h 100 μM VH298. **C)** ATAC-qPCR analysis in A549 cells cultured at 21% oxygen and with treated with 24h DMSO (control), 24h hypoxia and 24h 100 μM VH298. Graphs show mean (*N3*) ± SEM, \**P*< 0.05, \*\**P*< 0.01, \*\*\**P*< 0.001.

An open region of the *ACTB* promoter, which was unchanged in response to hypoxia and VH298 treatment in ATAC-seq analysis, was also analysed via ATAC-qPCR as a control (Supplementary Figure S4A-C). Immunoblotting for HIF-1α was also performed in A549 cells treated with 24h hypoxia or 24h, 100μM VH298 to confirm the expected hypoxia responsiveness/HIF-1α stabilisation in this cell line (Supplementary Figure S4D).

**Supplementary Figure S4.**
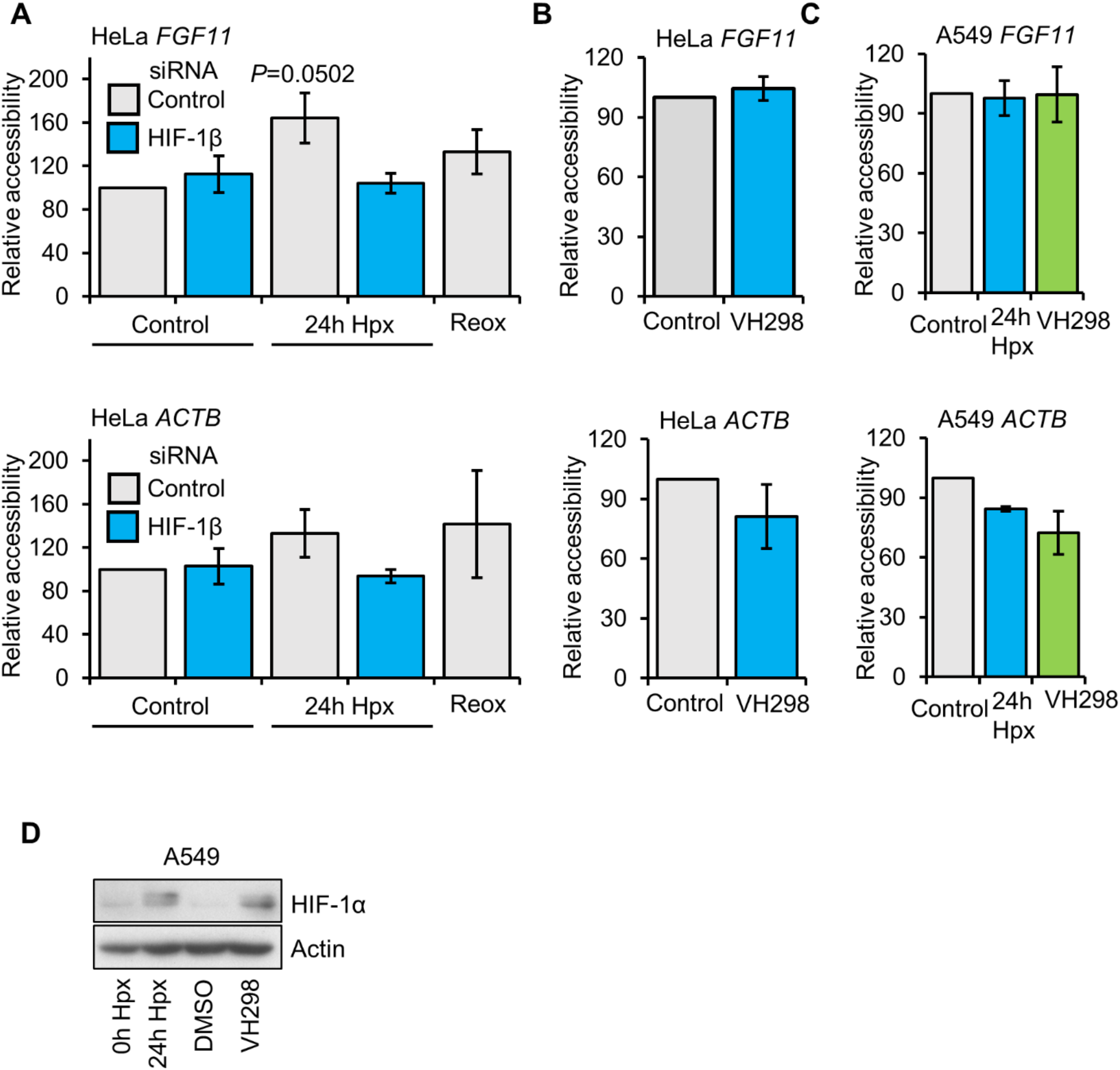
Validation of accessibility changes additional data. **A)** ATAC-qPCR analysis in HeLa cells cultured at 21% oxygen, transfected with control siRNA or HIF-1β siRNA, and exposed to 0h (control), 24h 1% oxygen (hypoxia (Hpx)) and 24h hypoxia followed by 1h at 21% oxygen (reoxygenation). **B)** ATAC-qPCR analysis in HeLa cells cultured at 21% oxygen and treated with 24h DMSO (control) or 24h 100 μM VH298. **C)** ATAC-qPCR analysis in A549 cells cultured at 21% oxygen and with treated with 24h DMSO (control), 24h hypoxia and 24h 100 μM VH298. Graphs show mean (*N3*) ± SEM, \**P*< 0.05, \*\**P*< 0.01, \*\*\**P*< 0.001. **D)** Immunoblot of the indicated proteins in A549 cells cultured at 21% oxygen and with treated 24h hypoxia, 24h 100 μM VH298 and 24h DMSO.

### Mechanistic insight into hypoxia inducible changes in chromatin accessibility

To gain mechanistic insight into hypoxia/HIF driven changes in chromatin accessibility, we measured the percentage of HIF binding sites present at VH298 and hypoxia DORs using HeLa HIF subunit ChIP-seq data (Figure 6A). HIF-1α, HIF-1β and HIF-2α binding sites are enriched at VH298 and 24h hypoxia upregulated DORs but not at downregulated DORs (Figure 6A). This indicates a role of direct HIF binding in hypoxia/VH298 induced increases in chromatin accessibility complemented by HIF indirect/independent changes. Next, we analysed HIF subunit binding sites at promoter, gene body and intergenic hypoxia or VH298 upregulated DORs, finding HIF subunit binding sites show the strongest preference for promoter DORs (Figure 6B, Supplementary Figure S5A). HIF binding sites are also more strongly enriched at DORs upregulated in response to both hypoxia and VH298 compared to hypoxia unique and VH298 unique upregulated DORs, suggesting that hypoxia unique and VH298 unique DORs may involve more HIF indirect changes (Figure 6C, Supplementary Figure S5B). Statistically significant overlaps of HIF binding sites are present at reoxygenation sensitive and HIF-1β dependent hypoxia upregulated DORs, but not at reoxygenation insensitive and HIF-1β independent hypoxia upregulated DORs (Figure 6D, Supplementary Figure S5C). Thus, HIF binding is a determinant of reoxygenation sensitivity and HIF-1β dependence regarding hypoxia upregulated DORs.

**Figure 6.**
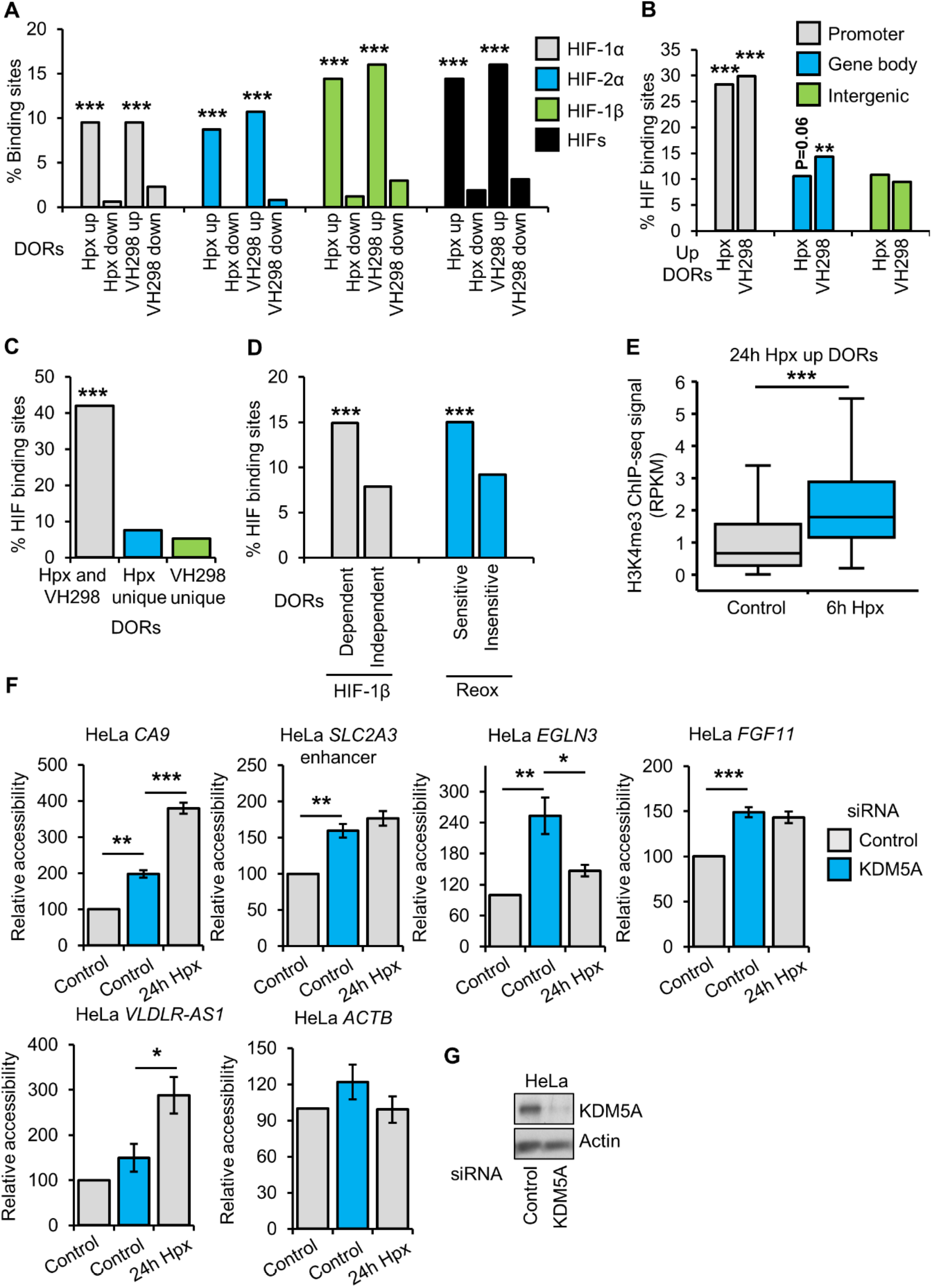
Mechanistic insight into hypoxia inducible changes in chromatin accessibility. **A)** Overlap of HIF subunit binding sites with 24h 1% oxygen (hypoxia (Hpx)) and 24h, 100 μM VH298 differentially open chromatin regions (DORs) identified by ATAC-seq in HeLa cells (\*\*\**P*< 0.001). **B)** Overlap analysis of HIF subunit binding sites with promoter, gene body and intergenic 24h hypoxia and VH298 upregulated DORs (** *P*< 0.01, \*\*\**P*< 0.001). **C)** Overlap of HIF subunit binding sites with both VH298 and 24h hypoxia upregulated DORs, 24h hypoxia unique DORs and VH298 unique DORs (\*\*\**P*< 0.001). **D)** Overlap of HIF subunit binding sites with HIF-1β dependent and independent 24h hypoxia upregulated DORs, and reoxygenation sensitive and insensitive 24h hypoxia upregulated DORs (\*\*\**P*< 0.001). **E)** Box plot of H3K4me3 ChIP-seq signal (RPKM) in HeLa cells exposed to 0h (control) and 6h hypxoia at 24h hypoxia upregulated DORs (centre +/- 1kb) (\*\*\**P* < 0.001). **F)** ATAC-qPCR analysis in HeLa cells cultured at 21% oxygen, transfected with control or KDM5A siRNA, and exposed to 0h (control) or 24h hypoxia. Graphs show mean (*N3*) ± SEM, \**P*< 0.05, \*\**P*< 0.01, \*\*\**P*< 0.001. **G)** Immunoblot analysis of the indicated proteins in HeLa cells transfected with control or KDM5A siRNA.

To investigate a potential role of chromatin remodellers in hypoxia induced chromatin accessibility changes, we performed overlap analysis of VH298 and 24h hypoxia DORs with publically available genome wide occupancy data in HeLa cells for members of the SWI/SNF complex (BRG1, BRG1-Associated Factor 155 (BAF155), BAF47, BAF170) and Sucrose Nonfermenting Protein 2 Homolog (SNF2H) (Supplementary Figure S6A). While there were no statistically significant overlaps, SWI/SNF binding sites were more prevalent at VH298 and 24h hypoxia upregulated DORs compared to downregulated DORs. To test if SWI/SNF is required for hypoxia inducible increases in chromatin accessibility, we depleted BAF155, a core subunit of SWI/SNF complexes, with siRNA, and measured chromatin accessibility changes at the previously validated sites (Figure 5) by ATAC-qPCR (Supplementary Figure S6B). At all 6 hypoxia upregulated DORs analysed, BAF155 depletion in hypoxia did not affect chromatin accessibility in hypoxia, demonstrating BAF155 is not required for hypoxia mediated increases in chromatin accessibility, at least the sites studied (Supplementary Figure S6B). An open region of the *ACTB* promoter, unaffected by hypoxia and VH298 treatment by ATAC-seq, was used as a control region (Supplementary Figure S6B). Immunoblot analysis confirmed effective depletion of BAF155 with siRNA treatment (Supplementary Figure S6C). As there is publically available HeLa cell ChIP-seq data for p300, a known HIF-α co-activator (23), we compared hypoxia/VH298 DORs with p300 binding sites (Supplementary Figure S6D). However, as with the chromatin remodeller overlaps, we found no statistically significant overrepresentation of p300 binding sites (Supplementary Figure S6D). A limitation of this analysis is that the p300 and chromatin remodeller data is in normoxic cells.

Changes to H3K4me3 in hypoxia correlate with changes in gene expression (12, 24) and have been linked to coordination of hypoxia induced transcriptional changes (12). Analysis of publically available H3K4me3 ChIP-seq data finds that in hypoxia, H3K4me3 is enriched at 24h hypoxia upregulated DORs (Figure 6E). As Lysine Demethylase 5A (KDM5A) depletion in normoxia has been shown to mimic hypoxia induced H3K4me3 and gene expression changes, we tested whether KDM5A depletion mimics hypoxia induced changes in chromatin accessibility (Figure 6F). ATAC-qPCR reveals that KDM5A depletion in normoxic HeLa cells increases chromatin accessibility at 4/5 hypoxia upregulated DORs analysed (Figure 6F). Immunoblot analysis confirmed effective depletion of KDM5A with siRNA treatment (Figure S6). Taken together, these data suggest an intricate crosstalk between HIF and KDM5A in the control of hypoxia-induced chromatin structure changes.

**Supplementary Figure S5.**
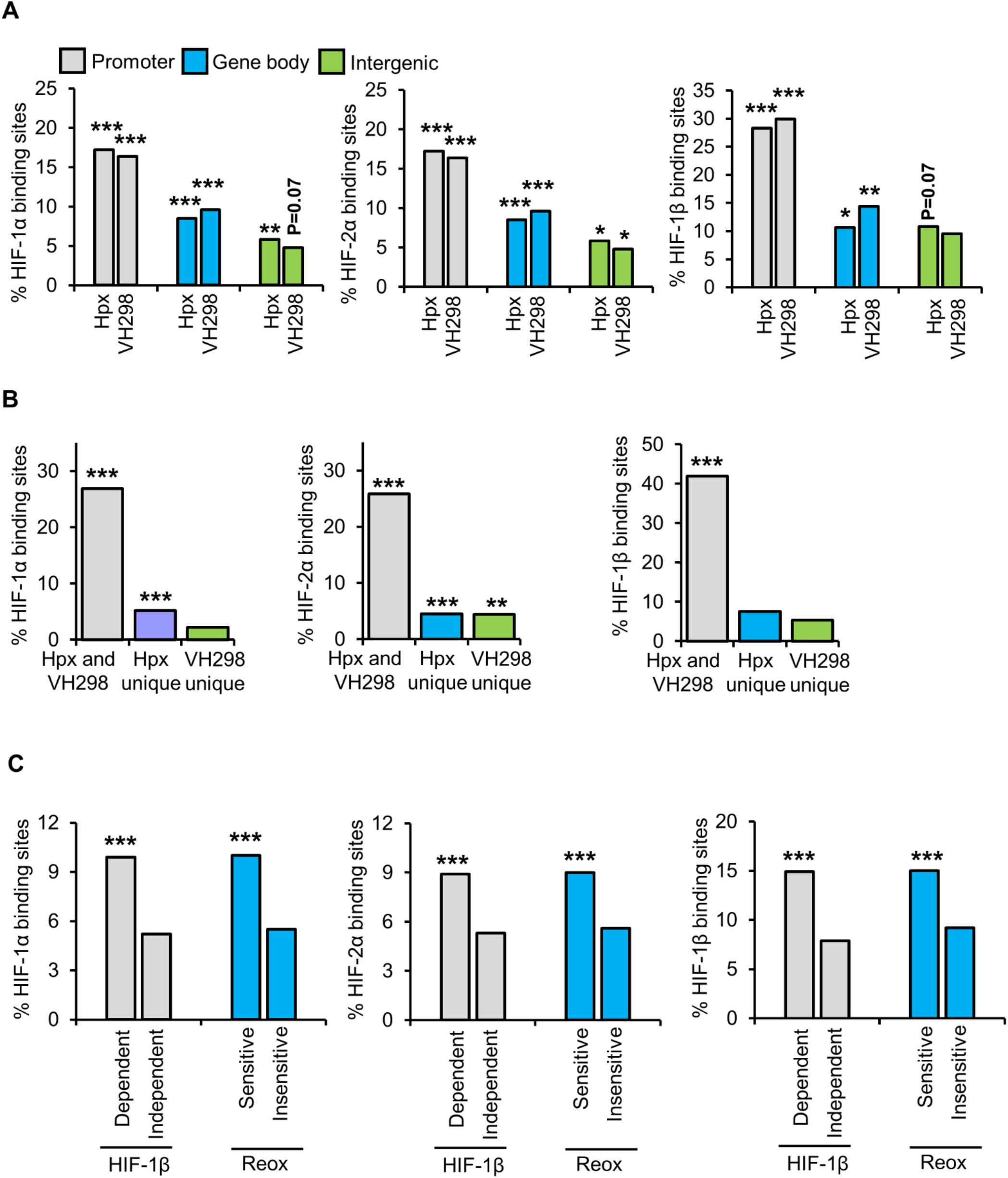
Additional HIF subunit binding analysis. **A)** Overlap of HIF subunit binding sites with promoter, gene body and intergenic 24h 1% oxygen (hypoxia (Hpx)) and 24h, 100 μM VH298 differentially open chromatin regions (DORs) identified by ATAC-seq in HeLa cells (\**P*< 0.05, \*\**P*< 0.01, \*\*\**P*< 0.001). **B)** Overlap analysis of HIF subunit binding sites with both VH298 and 24h hypoxia upregulated DORs, 24h hypoxia unique DORs and VH298 unique DORs (\*\**P*< 0.01, \*\*\**P*< 0.001). Overlap analysis of HIF subunit binding sites with HIF-1β dependent and independent 24h hypoxia upregulated DORs, and reoxygenation sensitive and insensitive 24h hypoxia upregulated DORs (\*\*\**P*< 0.001).

**Supplementary Figure S6.**
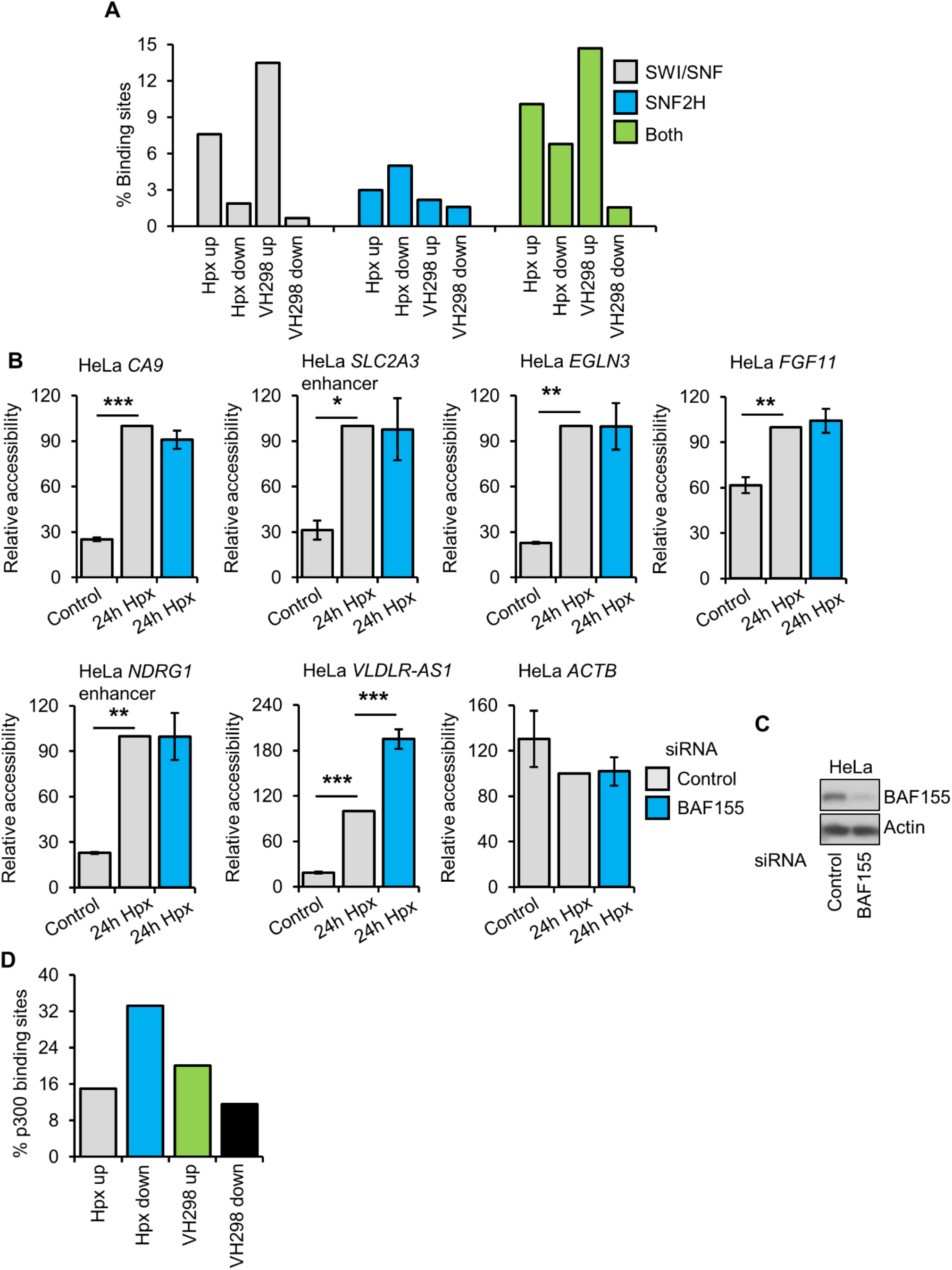
Chromatin remodeller and p300 analysis. **A)** Overlap of SWI/SNF subunit binding sites (BRG1, BAF47, BAF155 and BAF170) and SNF2H binding sites with 24h 1% oxygen (hypoxia (Hpx)) and 100 μM VH298 differentially open chromatin regions (DORs) identified by ATAC-seq in HeLa cells. **B)** ATAC-qPCR analysis in HeLa cells cultured at 21% oxygen, transfected with control or BAF155 siRNA, and exposed to 0h (control) or 24h hypoxia. Graphs show mean (*N3*) ± SEM, \**P*< 0.05, \*\**P*< 0.01, \*\*\**P*< 0.001. **C)** Immunoblot of the indicated proteins in HeLa cells transfected with control or BAF155 siRNA. **D)** Overlap of p300 binding sites with 24h hypoxia and VH298 upregulated DORs.

## Discussion

ATAC-seq is utilised here to measure the chromatin accessibility landscape to response to hypoxia. Our findings that low oxygen triggers loci specific changes in chromatin accessibility agrees with previous ATAC-seq studies in other cell lines exposed to hypoxia (16–19). We characterised genomic loci with differential accessibility in hypoxia, via integrative analysis with RNA-seq and publically available databases. This analysis reveals that changes in chromatin accessibility in hypoxia are associated with hypoxia responsive genes, both within genes and genes promoters, and at distal regulatory elements. Genes with differential accessibility include the core, well-characterised hypoxia responsive genes, including *CA9, EGLN3, SLC2A3* and *NDRG1*. Interestingly, the loci with one the biggest hypoxia induced change in accessibility is on *VLDLR-AS1*, an anti-sense transcript of the hypoxia inducible HIF target gene, *VLDLR* (*25, 26*). *VLDLR-AS1* gene expression was also elevated in hypoxia in our RNA-seq analysis in HeLa and A549 cells. This may represent a feedback loop to reduce *VLDLR* expression levels after prolonged exposure to hypoxia, similar to *HIF1A* and *HIF1A-AS2* (*27*). However, further studies will be needed to confirm this hypothesis.

HIF is central to transcriptional responses in hypoxia (2–5). Motif enrichment analysis identifies HIF subunit motifs are enrichment, specifically at sites with increased accessibility in hypoxia. By combining siRNA depletion of HIF-1β in cells exposed to normal oxygen and low oxygen, with ATAC-seq, we determine the dependence of HIF on hypoxia induced chromatin accessibility variations. Most sites where shown to require HIF for hypoxia driven alterations to accessibility, with a stronger dependence for upregulated accessibility loci compared to downregulated accessibility loci. Many cellular responses to low oxygen are highly dynamic and reverted upon reoxygenation, including HIF pathway activation, due to oxygen sensing via oxygen dependent enzymes (1, 8). Our analysis shows that the majority of accessibility changes in hypoxia are restored to normoxic levels following a short period of reoxygenation, demonstrating rapid and dynamic oxygen sensitivity, which parallels that of the HIF pathway.

Whilst the central oxygen-sensing pathway in metazoans is the VHL-PHD-HIF axis, impairment of other oxygen sensitive enzymes in low oxygen, and non-HIF targets PHD enzymes, can trigger other oxygen sensitive changes (14). This includes changes to DNA and histone methylation (28). In an attempt to delineate HIF stabilisation in hypoxia, from other effects caused by inhibition of cellular oxygen sensors, we performed ATAC-seq in cells treated with a chemical stabiliser of HIF-α, called VH298. This compounds works via blocking the hydroxylated HIF-α binding pocket of VHL, thus stabilising HIF-α and partially mimicking HIF mediated responses, without inhibiting oxygen sensors (20). VH298 induces fewer changes in chromatin accessibility than hypoxia, and the changes induced by VH298 are also associated with hypoxia responsive genes and display HIF motif enrichment. 20% of hypoxia upregulated chromatin accessibility sites are also increased in response to VH928. This analysis demonstrates that HIF stabilisation is sufficient to trigger increases in chromatin accessibility, which partially mimic those driven by hypoxia. A similar trend was observed when previously elucidating proteome wide and transcriptome wide changes in response to hypoxia and VH298 (21, 22). There was strikingly little overlap between reduced accessibility loci in hypoxia and VH298. A limitation of the VH298 ATAC-seq experiment is we only use 1 time-point of VH298 and it is known that hypoxia and VH298 have different dynamics regarding HIF stabilisation and activation of HIF target genes (20–22). Future work using multiple timpoints of VH298 and hypoxia will help distinguish change in accessibility driven solely by HIF stabilisation and those which require additional oxygen sensing mechanisms.

ATAC-seq findings are confirmed with ATAC-qPCR validation at a subset of hypoxia inducible genes with increased chromatin accessibility. Repeated analysis in second human cancer cell line, A549, uncovers cell type specific responses, with 2 out the 6 sites studied (*CA9 and FGF11 promoters*) only displaying accessibility sensitivity to hypoxia and VH298 in HeLa cells. Future investigation into chromatin responses across multiple cell types will help define cell conversed/specific responses as has been done previously for transcriptome response to hypoxia (29).

Comparison to pan genomic HIF binding sites unveils strong enrichment of HIF-1α, HIF2-α and HIF-1β at loci with increased accessibility in response to hypoxia and VH298 treatment. HIF binding is favoured at promoter hypoxia inducible accessible sites over gene body and intergenic loci and is HIF is also a determinant of HIF dependence and reoxygenation sensitive of hypoxia inducible accessible sites. Taken together, these data support a model whereby hypoxia inducible increases in chromatin accessibility are mostly HIF dependent, and consist of both direct local HIF binding and HIF indirect mechanisms, with a small contribution from HIF independent mechanisms. Conversely, around half of the loci with hypoxia-repressed accessibility are HIF independent and most of the HIF dependent sites are regulated indirect of local HIF binding.

Chromatin remodellers regulate cellular responses to hypoxia (30–32).We conducted preliminary investigation into the role played by chromatin remodellers in hypoxia and VH298 mediated accessibility changes by correlating hypoxia and VH298 responsive loci with pan genomic binding sites for normoxic SWI/SNF members and SNF2H binding sites in HeLa. No statistically significant correlations occur with this analysis, although it is limited by lack chromatin remodeller pan genomic occupancy studies in hypoxia. There is greater proportion of SWI/SNF binding sites at hypoxia and VH298 upregulated accessibility sites compared to downregulated accessibility sites. However, ATAC-qPCR analysis finds that a core subunit of the SWI/SNF complex, BAF155, is not required for hypoxia induced accessibility changes at the set of loci we validated.

Histone methylation modifications are sensitive to hypoxia (28) and KDM5A is reported cellular oxygen sensor that regulates H3K4me3 in hypoxia (12). We show that hypoxia induced upregulation of H3K4me3 is enriched at hypoxia upregulated accessibility sites. Furthermore, depletion KDM5A in normoxia, increases chromatin accessibility at some of validated hypoxia upregulated genomic loci. Thus, KDM5A may play a part in hypoxic regulation of chromatin accessibility. The histone methyltransferase SET1B was recently found to function as a HIF-1α coactivator, which is also required for H3K4me3 changes in hypoxia (24). As such, it will be important to elucidate the potential role of SET1B in hypoxia driven changes to the chromatin accessibility landscape. Histone acetylation and DNA methylation can influence chromatin accessibility, however these modifications and there effectors are not studied here.

Overall, our study provides the evidence for hypoxia induced chromatin structure changes that are extremely sensitive to oxygen, and primarily involve HIF. Further studies are needed to elucidate potential contributions and mechanisms for chromatin modifying enzymes and establish if chromatin accessibility changes in hypoxia and required for the hypoxia transcriptional response.

## Material and Methods

### Cell culture

Human cervix carcinoma HeLa and human lung carcinoma A549 cell lines were obtained from the American Type Culture Collection (ATCC) (Manassas, VA, USA) and maintained in Dulbecco’s modified Eagle’s medium (DMEM) (Gibco/ThermoFisher, Paisley, UK) supplemented with 10% v/v foetal bovine serum (FBS) (Gibco/ThermoFisher, Paisley, UK), 2 mM L-glutamine (Lonza, Slough, UK), 100 units/mL penicillin (Lonza, Slough, UK) and 100 μg/mL streptomycin (Lonza, Slough, UK) at 5% CO_2_ and 37 °C. Cell lines were cultures for no more than 30 passages and routinely tested for mycoplasma contamination using MycoAlert Mycoplasma Detection Kit (Lonza, Slough, UK).

### Treatments

Hypoxia treatments were performed by incubating cells in an InvivO2 300 Hypoxia Workstation (Baker Ruskinn, Bridgend, Wales) at 1% O_2_, 5% CO_2_ and 37 °C. Lysis of hypoxia treated cells was carried inside the hypoxia workstation to avoid reoxygenation. Reoxygenation treatments were performed by incubating cells for 24h in hypoxia followed by 1h incubation at 21% O_2_, 5% CO_2_ and 37°C. VHL binding to HIF-1/2α was inhibited by treating cells with 100 μM VH298 (Sigma, Gillingham, UK) for 24h and DMSO was used as vehicle control (Sigma, Gillingham, UK).

### siRNA transfections

Cells were transfected with 27 nM of small interfering RNA (siRNA) oligonucleotides (Eurofins, Ebersberg, Germany) for 48h using Interferin (Polyplus, Illkirch, France) transfection reagent according to manufacturer’s instructions. The following siRNA were used: Control CAGUCGCGUUUGCGACUGG, HIF-1β GGUCAGCAGUCUUCCAUGA, KDM5A GAAGAAUUCUAGCCAUACA.

### Immunoblotting

Cells were lysed in RIPA buffer (50 mM Tris-HCl, pH 8, 150 mM NaCl, 1% v/v NP-40, 0.25% w/v Na-deoxycholate, 0.1% w/v SDS, 10 mM NaF, 2 mM Na_3_VO_4_ and 1 tablet/10 mL, Complete, Mini, EDTA-free protease inhibitor (Roche, Welwyn Garden city, UK)). Samples were incubated for 10mins on ice, centrifuged at 13000rpm, 10min and 4°C, and supernatants (RIPA soluble protein lysates) were collected. Standard SDS-PAGE and immunoblotting protocols were performed with 20 μg of protein per lane loaded on SDS-PAGE gels. The following primary antibodies were used for immunoblotting: HIF-1α (610958, BD Biosciences (Workingham, UK)), HIF-1β (3718, CST ((Leiden, Holland)), Actin (60009-1, Proteintech (Manchester, UK)), BAF155 (11956, CST ((Leiden, Holland)), KDM5A (3876, CST ((Leiden, Holland)). 3 biological replicates were analysed per condition. Immunoblot figures are from 1 biological replicate, which is representative of all replicates.

### ATAC-seq

Assay for Transposase-Accessible Chromatin using sequencing (ATAC-seq) was performed using the following protocol adapted from (33, 34). Cells were washed directly on cell culture plates with in 2 mL DPBS (Gibco/ThermoFisher, Paisley, UK) and 1 mL of resuspension buffer (10 mM Tris-HCl, pH 7.5, 10 mM NaCl, 3 mM MgCl_2_). Lysis buffer (0.1% v/v NP-40, 0.1% v/v Tween-20, 0.1mg/mL Digitonin ((Promega, Southampton, UK) in resuspension buffer) was added at a volume resulting in a cell concentration of 1000 cells/uL followed by gentle scraping with a cell scraper and transfer of the cell suspension to 1.5 mL Eppendorf tubes. Samples were incubated on ice for 3min. 1 mL of wash buffer (0.1% v/v Tween-20 in resuspension buffer) was added and mixing was performed by inverting tubes 3 times. Samples were centrifuged at 1000g, 10min and 4°C. The supernatant was discarded and the pellet (cell nuclei) was resuspended in 50 uL transposition mix (50% v/v 2X Tagment DNA (TD) Buffer (Illumina, Cambridge, UK), 32% v/v PBS, 0.5μL final 0.1% v/v Tween-20, 0.1 mg/mL Digitonin (Promega, Southampton, UK), 5% v/v TDE1 Tagment DNA Enzyme (Illumina, Cambridge, UK) in nuclease free water (Sigma, Gillingham, UK)) by gentle pipetting up and down 6 times. Transposition (tagmentation) reaction was performed by incubating samples at 1000rpm, 30min and 37°C on a thermomixer. DNA was purified using the MinElute PCR Purification Kit (Qiagen, Manchester, UK) according to manufacturer’s instructions, with DNA eluted in 10uL Elution buffer from the kit. Tagmented DNA was amplified by PCR in the following reaction mix; 10 uL DNA, 10 uL nuclease free water (Sigma, Gillingham, UK) 2.5 uL of 25 μM forward primer (Nextera/Illumina i5 adaptors (Illumina, Cambridge, UK)), 2.5 uL of 25 μM reverse primer (Nextera/Illumina i7 adaptors (Illumina, Cambridge, UK)) and 25μl NEBNext^®^ Ultra™ II Q5 Master Mix (NEB, Hertfordshire, UK), with the following cycling conditions; 5min 72°C, 30sec 98°C and 11 cycles of 10sec 98°C, 30sec 63°C and 1min 72°C. Double-sided magnetic bead based DNA purification (to remove primer dimers and large >1,000 bp fragments) was performed using Agencourt AMPure XP beads (Beckman Coulter, High Wycombe, UK). DNA was quality controlled using an Agilent 2100 Bioanalyzer (Agilent, Stockport, UK), multiplexed, size selected (170-650 bp) using a Pipin prep (Sage Science, Beverly, MA, USA) and sequenced using S1 chemistry (paired-end, 2×50 bp sequencing) on a Novaseq sequencer (Illumina, Cambridge, UK). 2 biological replicates were analysed per condition.

### ATAC-seq data analysis

Reads in fastq files were trimmed for adaptors using Cutadapt (ref) and low quality score using Sickle. Reads were aligned to the human genome version hg38 (UCSC) using Bowtie2 (35), sorted and indexed binary alignment mapped (bam) files with mitochondrial reads removed were generated using Samtools (36). Bam files were filtered to keep ‘only properly paired reads’ following ENCODE guidelines using Samtools (36). PCR duplicates were removed from bam files using Picard. Number of reads in bam files and their fragment length distribution was determined using Samtools (36). Open chromatin regions (ORs) for each biological replicate were identified using MACS2 (37) (--nomodel --shift −100 --extsize 200 -q 0.01) and filtered to remove ENCODE DAC hg38 blacklisted regions and regions with an FDR < 1×10^-15^ using GenomicRanges (38) and ChIPpeakAnno (39). ORs for each replicate within a condition were overlapped using ChIPpeakAnno (39) and regions not present within the overlap were excluded. Library sized normalised (reads per kb per million reads (RPKM)) bigwig files and metagene graphs and heatmaps were made using deepTools (40). Differential open chromatin regions (DORs) between 2 conditions were determined using DiffBind (41) (dba.count fragmentSize = 150), dba.normalize library=DBA_LIBSIZE_PEAKREADS, dba.analyze method = DBA_DESEQ2) with filtering for log2 fold change (-/+0.58) and FDR (0.05). PCA plots were generated DiffBind (41). Closest gene TSS to ORs/DORs and genomic annotation of ORs/DORs were identified using ChIPpeakAnno (39). Overlap of DORs with each other or other genomic intervals was performed using ChIPpeakAnno (39). Genomic annotations of DORs were assigned using ChIPseeker(42). Promoters were defined as TSS -/+ 3kb, gene bodies were defined as regions more than 3kb downstream of TSS and upstream of TES and all other regions were defined as intergenic. Genic regions were defined as promoter and gene body regions. Gene signature analysis was performed using the Molecular Signatures Database with hallmark gene sets (43, 44). Motif enrichment analysis was performed using HOMER with DORs set as foreground and all ORs set as background (45). Gene set enrichment analysis was performed using WebGestalt (46). Enhancer analysis was performed using the HACER database (47). Volcano plots were made using R Bioconductor package EnhancedVolcano. Coverage tracks were produced using IGV (48).

### ATAC-qPCR

Pre-multiplexed ATAC-seq DNA was diluted to 0.5 ng/uL and qPCR analysis of chromatin accessibility was performed by running 3uL of DNA on a Mx3005P qPCR platform (Stratagene/Agilent, Stockport, UK) with Brilliant II Sybr green reaction mix (Stratagene/Agilent, Stockport, UK) in a final reaction of 15 uLs. The following qPCR primers were used; CA9 F CAGACAAACCTGTGAGACTTT and R TACGTGCATTGGAAACGAG, PHD3 F TACAGGGTGTTTGGGTTTG and R ACGTAGCCCTGTCACTC, FGF11 F CAGACAGACAGACAGACAGATG and R CGCTAGCTTGGCGAGAG, VLDR AS1 F CAGTCCCAGTGTGCATATTT and R CCTCTGGGTGTTAGCATTTC, ULK1 F GGTGGCCCTTCCTTCTTA and R GCTGGACAGAACCACTCT, ACTB F GCGGTGCTAGGAACTCAAA and R TACTCAGTGGACAGACCCAA, NDRG1 enhancer F AGAAGGTGTGCGTGTTTAG and R GATGACTCCAGAAACCAAGAG, GLUT3 enhancer F CTTAGTTGTATCTGGGTGTGG and R GAGAGGAGCAATGTCTGATG. 3 biological replicates were analysed per condition.

### RNA-seq

RNA was extracted from HeLa and A549 cells using an RNeasy Mini Kit (Qiagen, Manchester, UK). RNA was quality controlled using an Agilent 2100 Bioanalyzer (Agilent, Stockport, UK). Dual-indexed, strand specific RNA-seq libraries were generated using NEBNext polyA selection and Ultra Directional RNA library preparation kits (NEB, Hertfordshire, UK), multiplexed, and sequenced (Paired-end, 2×150 bp sequencing) on a HiSeq 4000 sequencer (Illumina, Cambridge, UK). 3 biological replicates were analysed per condition.

### RNA-seq data analysis

Reads in fastq files were trimmed for adaptors using Cutadapt and low quality score using Sickle (23). Reads were aligned to the human genome version hg38 (GRCh38, Ensembl) using STAR (49) and the resulting binary alignment mapped (bam) files were indexed using Samtools (36). Read counts for each transcript (GRCh38, Ensembl) were generated using Subread (featureCounts) (50). Differential expression analysis were performed using R Bioconductor package DeSeq2 (51), with filtering for log2 fold change (-/+0.58) and FDR (0.05).

### Statistical analysis

For ATAC-qPCR analysis comparing 2 conditions, statistical significance was determined via Student’s t-test. For ATAC-qPCR analysis comparing more than 2 conditions, statistical significance was determined via one-way ANOVA with post-hoc Tukey test. For overlap of genes with differentially accessible chromatin regions identified by ATAC-seq, with genes with differential RNA expression, statistical significance was determined via Fisher’s exact test. For comparison of H3K4me3 ChIP signals, statistical significance was determined via Wilcoxon signed-rank test. For all other statistical analysis, default settings of the particular analysis tool were used. In all cases, \**P* < 0.05, \*\**P*< 0.01, \*\*\**P*< 0.001.

### Data mining of public available datasets

HeLa HIF and H3K4me3 ChIP-seq datasets (24)(GSE169040 and GSE159128) and HeLa ATAC-seq datasets (52, 53) (GSE121840 and GSE106145) were downloaded from the Gene Expression Omnibus (54). p300 (ENCFF631WOD), BRG1 (ENCFF216YDM), BAF47 (ENCFF572FHR), BAF155 (ENCFF492BST) and BAF170 (ENCFF253UAA) ChIP-seq datasets were downloaded from the ENOCDE portal (55). SNF2H ChIP-seq (56)(PRJEB8713) dataset was downloaded from the European Nucleotide Archive.

## Supplementary Datasets

**Supplementary dataset 1.** ATAC-seq quality control and open chromatin regions (ORs).

**Supplementary dataset 2.** ATAC-seq differential open chromatin region (DOR) analysis.

**Supplementary dataset 3.** RNA-seq differential expressed genes (DEGs).

**Supplementary dataset 4.** ATAC-seq and RNA-seq integrative analysis.

**Supplementary dataset 5.** ATAC-seq HIF-1β independent differential open chromatin regions (DORs).

**Supplementary dataset 6.** ATAC-seq reoxygenation sensitive/insensitive differential open chromatin regions (DORs).

**Supplementary dataset 7.** Hypoxia and VH298 ATAC-seq differential open chromatin regions (DORs) overlap analysis.

## Data availability

ATAC-seq (GSE186342 and GSE186123) and RNA-seq (GSE186370) data are deposited at the Gene Expression Omnibus (54).

## Acknowledgements

We would like to acknowledge the Centre for Genomic Research (CGR) at the University of Liverpool for quality control, multiplexing and sequencing of ATAC-seq samples, and quality control, library preparation, multiplexing and sequencing of RNA-seq samples.

## Competing interests

The authors declare no competing interests.

## Author contributions

M.B and S.R conceptualised and oversaw the project. M.B. performed all experiments and analysed the data unless otherwise indicated. J.F. performed VH298 ATAC-seq experiments. D.S. performed siRNA depletion immunoblotting. M.B. and S.R. wrote the manuscript. All authors have read and agreed to the published version of the manuscript.

## Notes

### Competing Interest Statement

The authors have declared no competing interest.

